# KNOWLEDGE GRAPH AIDS COMPREHENSIVE EXPLANATION OF DRUG TOXICITY

**DOI:** 10.1101/2022.10.07.511348

**Authors:** Yun Hao, Joseph D. Romano, Jason H. Moore

## Abstract

In computational toxicology, prediction of complex endpoints has always been challenging, as they often involve multiple distinct mechanisms. State-of-the-art models are either limited by low accuracy, or lack of interpretability due to their black-box nature. Here we introduce AIDTox, an interpretable deep learning model which incorporates curated knowledge of chemical-gene connections, gene-pathway annotations, and pathway hierarchy. AIDTox accurately predicts cytotoxicity outcomes in HepG2 and HEK293 cells. It also provides comprehensive explanations of cytotoxicity covering multiple aspects of drug activity including target interaction, metabolism, and elimination. In summary, AIDTox provides a computational framework for unveiling cellular mechanisms for complex toxicity endpoints.

## INTRODUCTION

*In silico* toxicity assessment is becoming a critical step in drug development owing to sustained advances of machine learning techniques^1^. A prevalent approach for toxicity prediction is quantitative structure-activity relationship (QSAR) modeling, which links a toxicity outcome to structural properties of compounds^2^. Coupled with millions of data points generated by large-scale toxicity testing^3, 4^, various QSAR models have been proposed to predict *in vitro* and *in vivo* endpoints^5–13^. While some have achieved decent predictive performance, almost none can overcome the trade-off between accuracy and interpretability^14^. Under state-of-the-art models, it remains challenging to explain the toxicity outcomes of a compound with cellular activities involving target proteins, specific pathways, and biological processes. This limitation has raised substantial concerns among experimental toxicologists, calling for prediction models that can provide insight into cellular mechanisms of toxicity.

Recent developments in visible neural networks (VNN) provide a solution to the issue^15–17^. In contrast to black-box neural networks, connections within VNNs are guided by curated knowledge from pathway ontologies. Specifically, the fully connected structure is replaced by an interpretable ontological hierarchy that connects input gene features to output response via hidden pathway modules. In a previous study, we developed a VNN model— named DTox—for predicting compound response to toxicity assays^18^. Importantly, DTox advances an innovative interpretation framework that identifies network paths connecting genes and pathways for explaining the toxicity outcome of compounds. We demonstrated that DTox can achieve the same level of predictive performance as state-of-the-art models with a significant improvement in interpretability, as the identified paths can be linked to well-established mechanisms of toxicity. However, one major limitation of DTox is that the input feature profile is derived from structure-based binding prediction models. While these models ensure the wide applicability of DTox, they are vulnerable to prediction errors and data scarcity, causing exclusion of certain genes from the feature space. To address this limitation, we used curated knowledge from the toxicology-focused graph knowledge base ComptoxAI^19^ to refine the input feature profile of DTox. ComptoxAI provides extensive profiling of chemical-gene connections across multiple gene categories, which can be used for feature profile construction. Hence, the resulting model—named AIDTox (Comptox**AI** + **DTox**)—contains many novel gene features with active roles in cellular mechanisms of toxicity, including tubulin proteins that regulate apoptosis signaling, cytochrome P450 enzymes that are mainly responsible for drug metabolism, and transporters that participate in drug elimination. We believe AIDTox will facilitate the generation of new hypotheses for mechanistic investigation. Our code can be accessed openly at https://github.com/yhao-compbio/AIDTox.

## METHODS

### Processing cell viability screening datasets for model training

The Tox21 datasets^3^ contain screening results indicating the response of *in vitro* toxicity assays to compounds of interest. We extracted active and inactive compounds from the screening results of each assay, then removed compounds with inconclusive or ambiguous results. We focused our analyses on two cytotoxicity assays: HEK293 and HepG2, for each of which at least 500 compounds are available for model training (**Table S1**).

### Extracting chemical-gene connections from ComptoxAI for model construction

ComptoxAI^19, 20^ is a comprehensive graph knowledge base that contains curated relationships between chemicals, genes, assays, and many other entities. It contains two types of relationships linking compound and gene nodes: physical binding (with protein product) and expression-alteration (up-regulation/down-regulation). We extracted both types for chemicals present in the Tox21 datasets. We also constructed a hybrid type by combining the two types of relationships. We used the extracted relationships to assemble binary compound-gene matrices as input feature profiles.

To perform dimensionality reduction, we implemented a ReliefF-based method—namely, MultiSURF^21^—for ranking genes by relevance to the assay outcome of interest. MultiSURF takes in a labeled feature dataset (assay outcome as the label), ranks all features based on differences among neighboring instances. Specifically, intraclass differences will contribute negatively to feature relevance while interclass differences will contribute positively. A benchmark study showed that MultiSURF outperformed other methods in detecting genotype-phenotype associations^21^. We selected the top 100, 200, 300, 400, and 500 ranked genes of each dataset to proceed with the following analysis. Model selection by training loss was carried out to identify the optimal gene feature space (**Table S2**).

### Constructing VNN with selected gene features and Reactome pathway hierarchy

DTox is a VNN model embedded with the Reactome pathway hierarchy that comprises root biological processes, child-parent pathway relations, and gene-pathway annotations (downloaded in Aug 2019)^22^. Each pathway is embedded as a module consisting of hidden neurons. AIDox inherits the basic structure of DTox while modifying input layers to consist of the gene features selected by MultiSURF. Guided by gene-pathway annotations, the gene features are connected to modules representing lowest-level pathways. These modules are then connected to deeper layers based on child-parent pathway relations until root biological processes are reached. Finally, the root biological processes are connected to an output layer representing the assay outcome. To trim the network’s scale and prevent overfitting, we made the root biological process a hyperparameter of AIDTox. We selected four processes (‘gene expression’, ‘immune system’, ‘metabolism’, and ‘signal transduction’) and all possible combinations among them for the tuning process (15 values in total). These four processes were selected due to their broad coverage and direct involvement in cellular mechanisms of toxicity.

AIDTox also inherits the hybrid loss function of DTox combing both root and auxiliary loss terms: 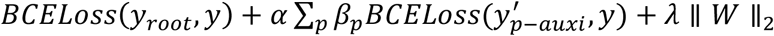. The first term, namely root loss, denotes the binary cross entropy of final output *y*_*root*_. The second term, namely auxiliary loss, denotes the binary cross entropy of the auxiliary scalar 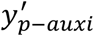 from each pathway module, with factor α (= 0.5) balancing the root and auxiliary terms, and factor *β*_*p*_ (computed as the inverse of pathway count within the corresponding hidden layer) normalizing terms across hidden layers. The third term denotes *L*_2_ regularization with coefficient *λ* (= 1e^−4^). The incorporation of auxiliary loss terms can prevent gradients from vanishing in the lower hierarchy and facilitates learning new patterns from individual pathways.

### Learning optimal AIDTox model for cytotoxicity prediction and interpretation

AIDTox adopts the same training scheme as DTox, including a training/testing/validation split by ratio of 7:1:2, optimization using the Adam algorithm^23^ with mini-batch size of 32, a learning rate of 0.001, early stopping criterion with “patience” of 20, and hyperparameter tuning by grid search. We evaluated the performance of the optimal AIDTox model on the held-out validation set (**Table S3**), and compared it with three existing models: (i.) the optimal DTox model, (ii.) an optimal QSAR model based on a random forest classifier (derived from tuning six hyperparameters with 2800 combinations in total, listed in **Table S4**), and (iii.) an optimal QSAR model based on a gradient boosting classifier (derived from tuning five hyperparameters with 3000 combinations in total, listed in **Table S4**). For the two QSAR models (ii. and iii.), we adopted 166-bit binary MACCS fingerprints to quantify structural properties of compounds, which covers most of the interesting physicochemical features for drug discovery^24^. We also implemented DTox’s model interpretation framework to identify paths from the VNN that can explain compound cytotoxicity (**Table S5**).

## RESULTS

### AIDTox employs curated chemical-gene connections to construct VNN

AIDTox predicts and explains compound response to toxicity assays with a knowledge-guided VNN. The knowledge incorporated in AIDTox comprises chemical-gene connections from ComptoxAI, as well as gene-pathway annotations and child-parent pathway relationships from Reactome (**Figure 1A**). Connections within the VNN are constrained to gene-pathway connections (input to first hidden layer) and child-parent pathway relations (after first hidden layer). In this study, we focused on two cell viability assays measuring compound cytotoxicity in HEK293 and HepG2 cells. Both datasets contain 1,367 compounds with connections in ComptoxAI, including 432 cytotoxic and 935 non-cytotoxic compounds in HEK293 cells, and 293 cytotoxic and 1,074 non-cytotoxic compounds in HepG2 cells. Among these compounds, 617 (45%) are connected to 991 genes with physical binding evidence, 1,237 (90%) are connected to 8,723 genes with expression-alteration evidence, and 1,367 (100%) are connected to 8,735 genes with both types of evidence. In all three cases, the number of gene features is much greater than the number of compound samples. To prevent overfitting, we reduced feature dimensions by selecting the top 100-500 predictive genes during model construction. In general, as more predictive genes were incorporated, we observed an improvement in the resulting model performance (decline in training loss) until 400 (**Figure S1**). Therefore, we proceeded with models built using the top 400 predictive genes.

**Figure 1.**
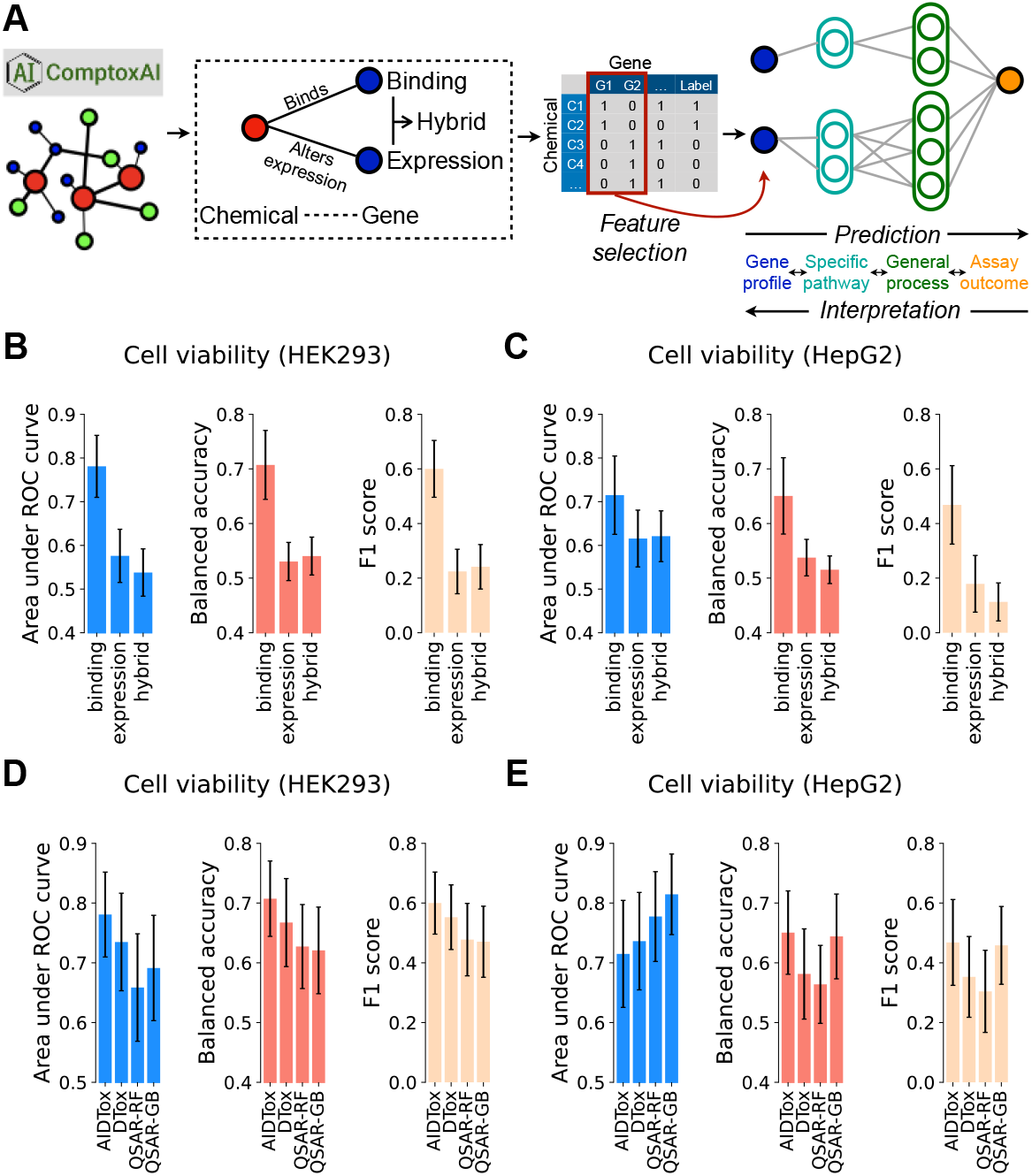
Incorporating curated chemical-gene connections into VNN for toxicity prediction with AIDTox. **(A)** Three types of chemical-gene connections (binding, expression, and hybrid) are extracted from ComptoxAI to construct the input feature profile of AIDTox. Feature selection is implemented to identify the top gene features predictive of the outcome of interest. The selected profile is fed into a VNN, whose structure is guided by Reactome pathway hierarchy. Specific pathways and general processes are coded as modules by hidden neurons. **(B&C)** Barplots showing the comparison of validation performance across three connection types in two cell viability datasets: HEK293 **(B)** and HepG2 **(C)**. Performance is measured by three metrics: area under ROC curve, balanced accuracy, and F1 score, with error bar showing the 95% confidence interval. **(D&E)** Barplots showing the comparison of validation performance across four models in two cell viability datasets: HEK293 **(D)** and HepG2 **(E)**. Three other models are considered: (i) our previous DTox model with inferred target profile as input, (ii) QSAR model by random forest with chemical fingerprint as input, and (iii) QSAR model by gradient boosting with chemical fingerprint as input.

### Chemical-gene binding connections result in the best performing models

Using held-out validation sets of HEK293 and HepG2 cell viability data, we first compared the performance of models derived from three distinct connection types. In both datasets, models derived from binding connections significantly outperform the other types (**Figure 1B&C; Table S2**), as we observed no overlaps between the 95% confidence intervals of compared metrics (AUROC, balanced accuracy, and F1-score) except for AUROC on the HepG2 viability dataset. We then compared the performance of AIDTox models derived from binding connections against three other classification algorithms, including our previous DTox model (derived from predicted compound-target interactions) and two QSAR models (derived from quantified structural properties). In both datasets, the 95% confidence intervals of performance metrics are highly overlapped among compared algorithms (**Figure 1D&E; Table S3**). Therefore, AIDTox achieved the same level of predictive performance as these well-established classification algorithms.

### AIDTox models benefit from a comprehensive gene feature space

A fundamental advantage of AIDTox over DTox comes from its enlarged gene feature space, providing an extensive profiling of the cellular activities of compounds. Overall, there is an increase from 361 genes in DTox to 991 genes in AIDTox, a 2.7-fold increase. Comparing the composition of gene features, we discovered that the increase in AIDTox is mainly driven by three target categories: ion channels, transporters, and others (**Figure 2A**). One example of the “others” category is the tubulin protein superfamily (**Figure 2C**), which polymerize into microtubules, a primary component of cytoskeleton involved in cell division. Due to the lack of binding data, tubulin proteins were not present in the DTox model. In ComptoxAI, the two families of tubulin proteins (α and β) are connected to antiparasitics (albendazole, and mebendazole) and antineoplastics (podofilox, vincristine sulfate, and docetaxel). Our AIDTox model further connects these drugs to HEK293 cytotoxicity via Rho GTPase signaling by effectors such as formins and IQGAPs, which have been shown to regulate apoptosis in HEK293 cells^25^.

**Figure 2.**
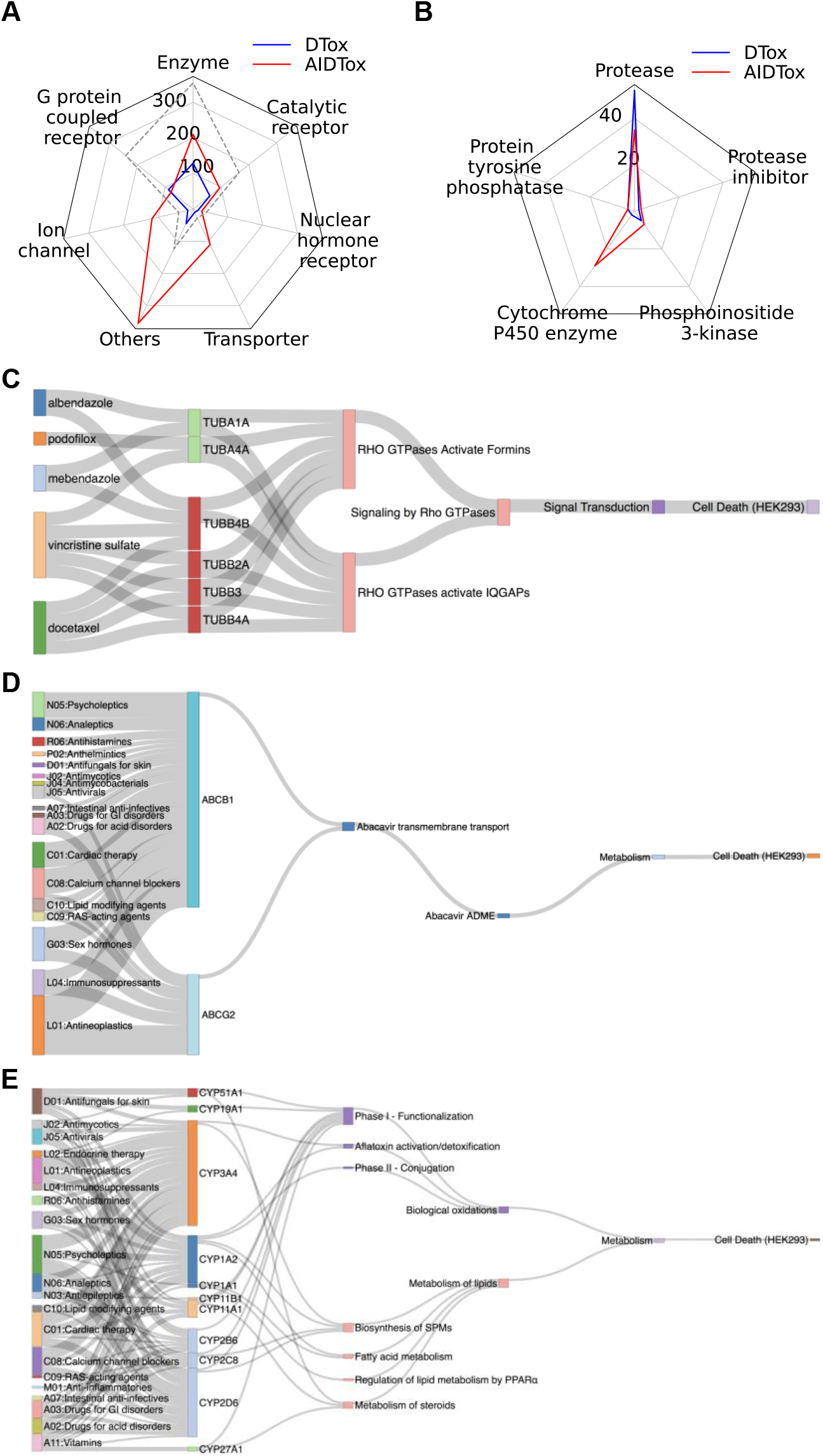
Comprehensive explanation of HEK293 cytotoxicity with new features in AIDTox. **(A)** Radar plot showing the comparison of gene category distributions among the features of DTox (blue solid line) and AIDTox (red solid line). The grey dashed line shows the hypothetical of a proportionate increase from DTox to AIDTox. **(B)** Radar plot showing the comparison of enzyme subcategory distributions among the features of DTox (blue solid line) and AIDTox (red solid line). **(C)** Sankey diagram showing the AIDTox explanation of HEK293 cytotoxicity for drugs targeting tubulin proteins. The paths (connecting drugs to HEK293 cell death) shown in the diagram are identified from the full network of VNN model by the AIDTox interpretation framework. Connections in the VNN are informed by ComptoxAI (chemical-gene) and Reactome (gene-pathway, child-parent pathway). Tubulin proteins are grouped and colored by the family. **(D)** Sankey diagram showing the AIDTox explanation of HEK293 cytotoxicity via ATP-binding cassette transporters (similar to C). Drugs are grouped and colored by the ATC subclass (first three digits). **(E)** Sankey diagram showing the AIDTox explanation of HEK293 cytotoxicity via cytochrome P450 enzymes (similar to C). Drugs are grouped and colored by the ATC subclass (first three digits). Cytochrome P450 enzymes are grouped and colored by the family. Metabolic pathways are grouped and colored by the general metabolic process they belong to (“Biological oxidations” or “Metabolism of lipids”).

### New features in AIDTox are essential in drug metabolism and elimination processes

We observed a disproportionate 11-fold increase (from 10 to 110) among transporters in the gene feature space of AIDTox. For instance, AIDTox connects drugs within nine ATC classes (18 subclasses) to HEK293 cytotoxicity via two new features: transporters *ABCB1* and *ABCG2* (**Figure 2D**). These two ATP-binding cassette transporters are mainly responsible for pumping drugs out of the cell, thus they play a critical role in drug elimination. Therefore, inhibition of transporter-mediated elimination by drugs may prolong their cytotoxic effect on HEK293 cells^26^.

While we did not observe a disproportionate increase among the general enzyme category, we did observe a 14.5-fold increase (from 2 to 29) among one subcategory: the cytochrome P450 enzymes (CYPs; **Figure 2B**). For instance, AIDTox connects drugs within nine ATC classes (20 subclasses) to HEK293 cytotoxicity via 20 CYPs of seven families (**Figure 2E**). These CYPs are the major enzymes involved in drug metabolism as they participate in metabolic pathways such as “biological oxidation” and “metabolism of lipids”. Meanwhile, CYP-mediated metabolic processes are the main source of reactive oxygen species, which can lead to cellular oxidative stress and trigger apoptosis/necrosis^27^.

## DISCUSSION

Rich knowledge in ComptoxAI provides an accurate and extensive profiling of chemical-gene connections. Here, we have explored the incorporation of these connections into knowledge-guided deep learning models for predicting and explaining compound cytotoxicity. We considered three types of connections for the task: physical binding, expression-alteration, and a hybrid type. Models derived from binding connections exhibit the best predictive performance, since physical binding is stronger evidence of direct interaction compared to expression alteration. In contrast to our previous work DTox, the new AIDTox model employs curated knowledge for generating input feature profile, thus is not prone to errors from binding prediction models. In addition, AlDTox is not restrained by the availability of compound-target binding data. The feature space of AIDTox comprises many genes with insufficient binding data, including ion channels, transporters, and CYP enzymes. These categories exhibit central roles in drug metabolism and elimination processes. Accordingly, they become an asset for prediction and explanation of complex toxicity outcomes, which may be trigged by multiple cellular mechanisms. For instance, AIDTox was able to connect dasatinib, a leukemia drug, to HEK293 cytotoxicity via multiple aspects of drug activity, including MAPK14-mediated regulation of apoptosis, CYP1A2/CYP3A4-mediated drug metabolism, and ABCB1/ABCG2-mediated drug elimination (**Figure S2**). It is also worth noticing the high interpretability of AIDTox is achieved without loss of accuracy. Therefore, we anticipate AIDTox to be applied in both prioritizing compounds for safety testing and generating new hypothesis for mechanistic investigation.

Despite these highlights, AIDTox did not significantly outperform existing methods in cytotoxicity prediction. This can be attributed to the sharp decline in sample size, as the profiling of chemical-gene connections by ComptoxAI only applies to well-studied compounds. A smaller sample size in turn leads to wider confidence intervals of model performance metrics. Consequently, even though we observed a moderate increase in the performance metrics, the increase is well within the 95% confidence interval. In the future, we expect the performance of AIDTox to be enhanced after chemical-gene connections become available for more compounds. We also acknowledge the recent development of graph neural network-based link prediction algorithms, which can fill in missing connections for under-studied compounds. Such algorithms may help us increase the sample size for AIDTox training and enhance model performance. In addition to increasing sample size, we think the incorporation of context-specific knowledge, such as cell line-specific chemical-gene connections, may further enhance the performance of AIDTox.

## AUTHOR CONTRIBUTIONS

J.H.M. and Y.H. conceived the AIDTox project. J.H.M., Y.H., and J.D.R. designed the AIDTox model and data analysis workflow. Y.H. and J.D.R. performed the analysis. J.H.M., Y.H., and J.D.R. interpreted the results and wrote the paper. All authors read and approved the final manuscript.

