## Supplementary material for "KNOWLEDGE GRAPH AIDS COMPREHENSIVE EXPLANATION OF DRUG TOXICITY": Figure S1, Figure S2

#### SUPPLMENTARY FIGURES

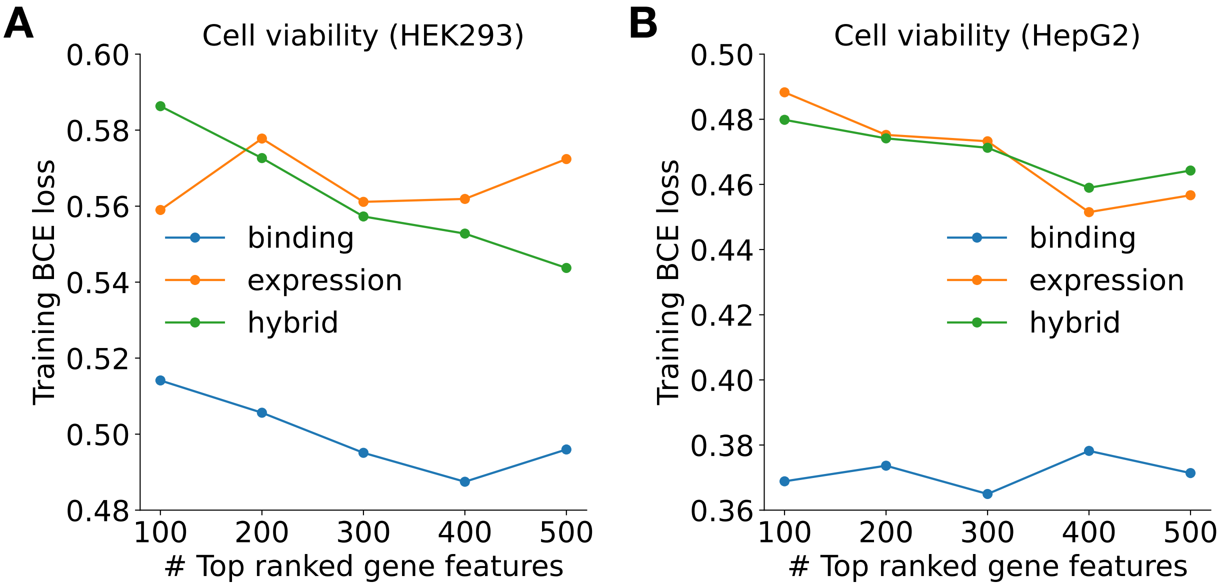


**Figure S1. Relationship between training performance and the number of top predictive gene features**

Line charts showing the relationship between training performance (y-axis) and the number of top predictive gene features (x-axis) in two datasets: HEK293 **(A)** and HepG2 **(B)** cell viability. Training performance is visualized by the binary cross entropy loss (BCE loss) for models derived from three types of chemical-gene connections: binding (blue line), expression (orange line), and hybrid (green line).

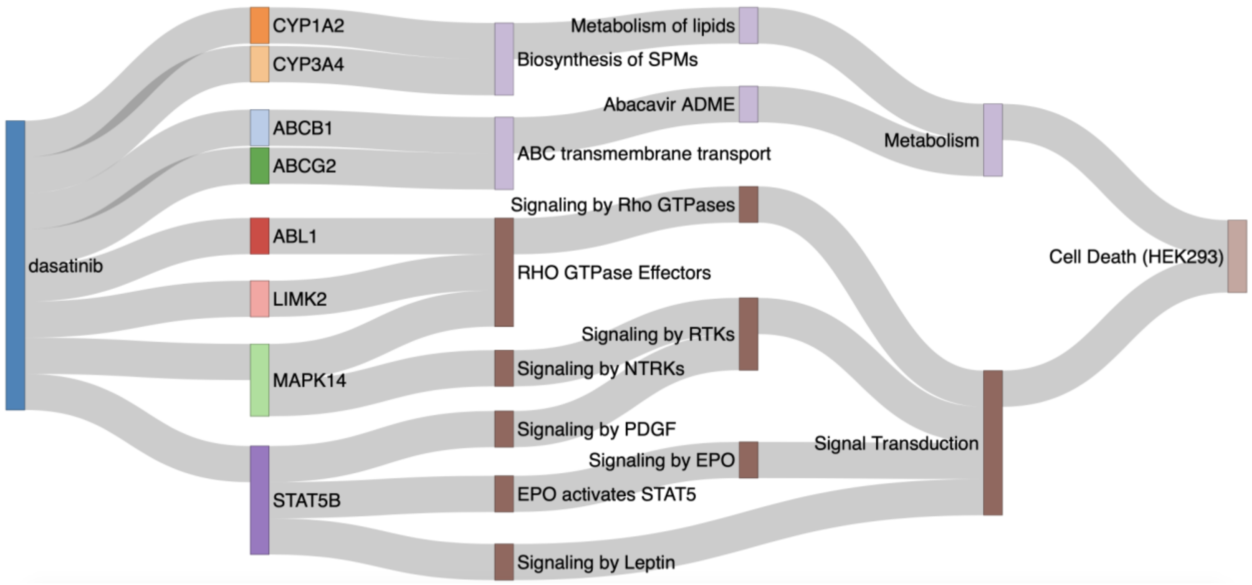


**Figure S2. AIDTox explanation for HEK293 cytotoxicity of dasatinib**

Sankey diagram showing the AIDTox explanation of HEK293 cytotoxicity for dasatinib, a drug used for leukemia treatment. The paths (connecting dasatinib to HEK293 cell death) shown in the diagram are identified from the full network of VNN model by the AIDTox interpretation framework. Connections in the VNN are informed by ComptoxAI (chemical-gene) and Reactome (gene-pathway, child-parent pathway). Pathways are grouped and colored by the general process they belong to ("Metabolism" or "Signal Transduction").

### SUPPLMENTARY TABLE TITLES

In the Excel file, we provide the following five supplementary tables:

**Table S1: Summary of two cell viability datasets used in the study**

**Table S2: Statistics of AIDTox models trained on cell viability datasets**

**Table S3: Evaluation of model performance on held-out validation cell viability datasets**

**Table S4: Hyperparameter tuning of classification algorithms**

**Table S5: VNN paths identified for active cytotoxic compounds of cell viability datasets**
